# The Cuban Human Brain Mapping Project population based normative EEG, MRI, and Cognition dataset

**DOI:** 10.1101/2020.07.08.194290

**Authors:** Pedro A. Valdes-Sosa, Lidice Galan, Jorge Bosch-Bayard, Maria L. Bringas Vega, Eduardo Aubert Vazquez, Samir Das, Trinidad Virues Alba, Cecile Madjar, Zia Mohades, Leigh C. MacIntyre, Christine Rogers, Shawn Brown, Lourdes Valdes Urrutia, Iris Rodriguez Gil, Alan C. Evans, Mitchell J. Valdes Sosa

**Affiliations:** The Clinical Hospital of Chengdu Brain Sciences, University of Electronic Sciences and Technology of China, Chengdu China; Cuban Center for Neuroscience, La Habana, Cuba; McGill Centre for Integrative Neurosciences MCIN. Ludmer Centre for Mental Health. Montreal Neurological Institute, McGill University, Montreal, Canada

**Author notes:** Co-first authors (Pedro A. Valdes-Sosa, Lidice Galan, Jorge Bosch-Bayard, Maria L. Bringas Vega, Eduardo Aubert Vazquez).

## Abstract

The Cuban Human Brain Mapping Project (CHBMP) repository is an open multimodal neuroimaging and cognitive dataset from 282 healthy participants (31.9 ± 9.3 years, age range 18–68 years). This dataset was acquired from 2004 to 2008 as a subset of a larger stratified random sample of 2,019 participants from La Lisa municipality in La Habana, Cuba. The exclusion included presence of disease or brain dysfunctions. The information made available for all participants comprises: high-density (64-120 channels) resting state electroencephalograms (EEG), magnetic resonance images (MRI), psychological tests (MMSE, Wechsler Adult Intelligence Scale -WAIS III, computerized reaction time tests using a go no-go paradigm), as well as general information (age, gender, education, ethnicity, handedness and weight). The EEG data contains recordings with at least 30 minutes duration including the following conditions: eyes closed, eyes open, hyperventilation and subsequent recovery. The MRI consisted in anatomical T1 and T2 as well as diffusion weighted (DWI) images acquired on a 1.5 Tesla system. The data is available for registered users on the LORIS database which is part of the MNI neuroinformatics ecosystem.

## Background & Summary

In the past decade, several healthy and patient neuroimaging databases of (ADNI, HCP, UK Biobank, CAMCAN, ABCD, PPMI) as well as consortia (ENIGMA) have been launched. They aim to accelerate discovery of insights into neurodevelopment and physiopathology, and to allow the identification of new biomarkers of disease with their integration into disease progression models. An essential ingredient, lacking many of these projects, is the inclusion of data from the electroencephalograms (EEG) as one of the most informative and direct measurement of brain activity--recordable at the same time scale of neural processes. At the same time, EEG is a cost effective and accessible neuroimaging modality that is applicable to underserved populations in all countries. It is now clear that EEG is a technique of choice for extensive population screening in any economic setting. This situation is being remedied with the collection and publication of open datasets such as ^1,2^.

This paper is an addition to the set of open multimodal datasets, having been originated in a Latin American middle-income country--the Cuban Human Brain Mapping Project (CHBMP) ^3^. This is a population-based, decades long, brain health data gathering effort in Havana Cuba. This still ongoing project is organized by the Cuban Ministry of Public Health (MINSAP) and coordinated by the Cuban Center for Neuroscience (CNEURO). The CHBMP focuses on the development of tools and health applications based on multimodal neuroimaging.

Making this dataset open is part of the Cuba-Canada-China (CCC) and the Global Brain Consortium (GBC strategy for open science, not only directed to integrate EEG neuroimaging as an essential component of future multimodal Neuroimaging projects, but also to serve as a “translational bridge” for resource limited scenarios in all countries. This has been made possible by integration of the CHBMP efforts into the MNI neuroinformatics ecosystem, based on the CBRAIN processing portal for the processing modules and the use of the LORIS database system^4^ for data storage and open access. This added value of the dataset is due to the health-oriented focus of the entire CHBMP which we now briefly summarize.

The emphasis of the CHBMP has been on quantitative evaluation, known as “qEEG” ^5^. qEEG has been shown to identify brain disorders in a wide variety of settings and therefore a candidate for use as a screening tool. In qEEG, the scalp recorded EEG log spectra of a proband are compared with normative spectra by means of a statistical parametric mapping procedure These normative spectra are obtained as age dependent means and standard deviations of the EEG log spectra of large sample of healthy participants. The age dependent norms are regressions of log-spectra over a wide age range. An EEG normative database is thus a prerequisite for qEEG.

The need for a Cuban normative database for qEEG ^6^ thus prompted the first wave of the CHBMP, initiated in 1988. This first database included 211 healthy persons from age 5 to 97 years. The participants were randomly selected from the Cuban population and screened by the Family Doctor system to include only healthy participants. This database was used to develop a high resolution qEEG validated by the Cuban Health system ^7,8^. It also promted the devlopment of qEEG for sources (qEEGt). Due to the unavailability of MRI in Cuba during the first wave CHBM, an “approximate qEEGt” was based on the average head model developed by the ICBM consortium ^9,10^. This first wave EEG dataset has being submitted separately ^11^ at Frontiers. Procedures to apply qEEG and qEEGt processing based on this dataset have been integrated into CBRAIN^12^.

At the end of the first wave of the CHBMP, it was recognized that qEEGt as in ^13^, based on individual MRI is important, not only to validate approximate qEEG, but also as a basis for multimodal neuroimaging studies of normal and pathological brain function. Thus, the need for a multimodal neuroimaging database was considered essential and planned for the time when MRI was available in Cuba. This was the motivation for the second wave of the CHBMP, between 2004 to 2008 as one of the projects for the National Program for Disability carried out at that time by the Cuban government. As in the first wave of the CHBMP the participants were recruited from the general population with a stratified randomized sampling of the population. This yielded 2,019 candidates, which were then screened, initially by the Family Nurses and Doctors, and later by extensive clinical, neurological, psychological, and neuroimaging evaluations in order to exclude participants with brain disorders, addictive habits or ill health. This resulted in a final sample of 282 “functionally healthy” participants. The recording protocol included high resolution EEG, T1 and T2 MRI, DWI, and psychological tests such as MMSE, Wechsler Adult Intelligence Scale (WAIS III), computerized reaction time, as well as the collection of blood samples for a genome wide association study (GWAS) to be described in a separate further publication.

It is to be noted that the third wave of the Cuban Human Brain Project was launched in 2019 and will be a large study of elderly subjects including a 10-year follow-up of the second wave CHBMP sample.

## Methods

### Participants

The Cuban Human Brain Mapping Project second-wave database contains neuroimaging, medical and cognitive data from, 282 “functionally healthy” participants from the general Cuban population between ages 18-68, (31.9 ± 9.30), comprising 87 (36.5±10.43) females and 195 (29.9 ±7.97) males. Details of the demographical variable frequency of gender, self-referenced handedness and educational level are presented in Table I.

**Table I:**
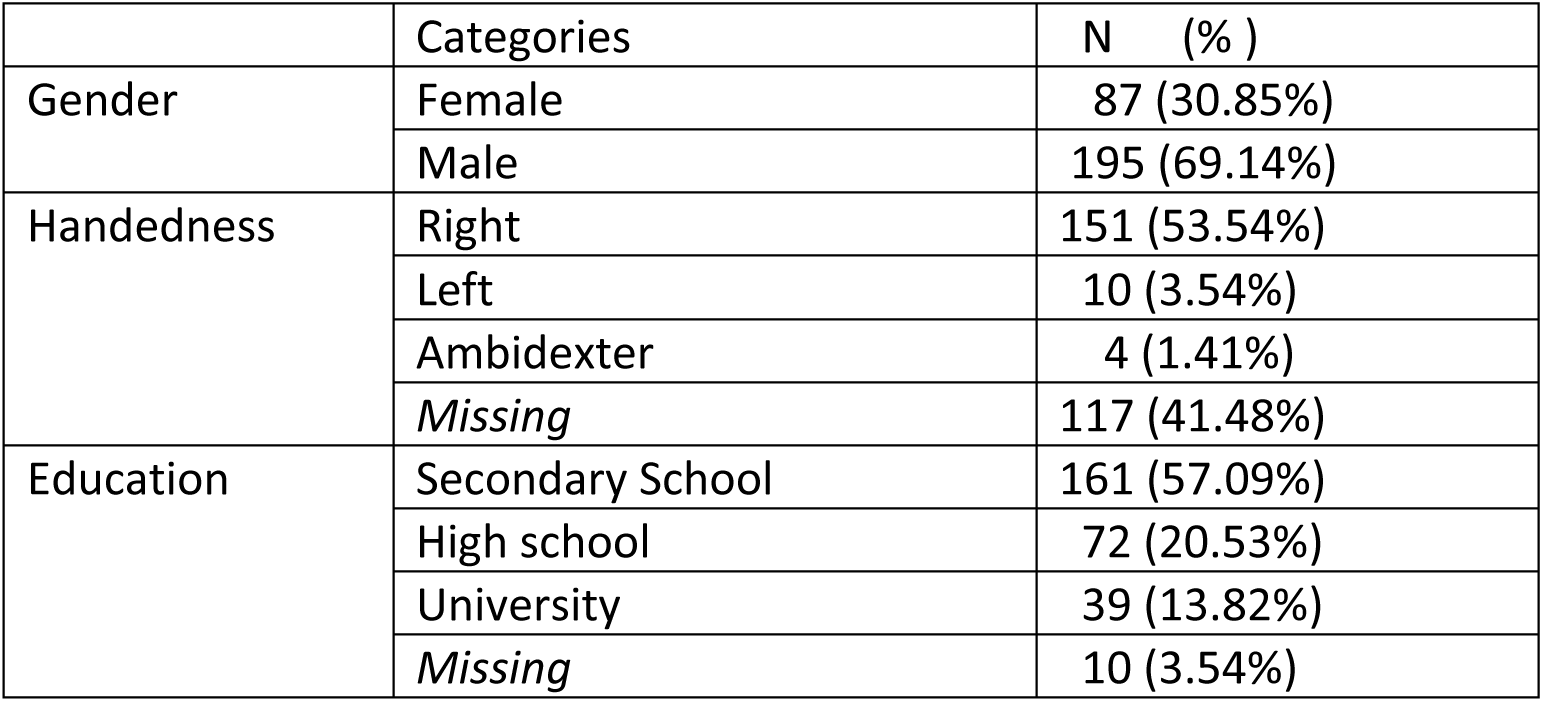
Demographic description of the sample.

### Ethics-

This study was carried out in accordance with the Declaration of Helsinki ^14^. The experimental protocols were approved by the Ethical Review Committee of the Cuban Center for Neuroscience using the guidelines of the Public Health (MINSAP from Spanish) and the Science, Technology and Environment Ministries of the Republic of Cuba (CITMA from Spanish). Participants included in the study signed a written consent after a house visit by the Primary Health care nurse who gave a verbal and written explanation of the purpose, risks and benefits of the study. Participants also were informed about the confidentiality of their personal information as well as full access to the best diagnostic and therapeutic procedures in case any kind of illness were detected that might prevent them to form part of the normative dataset. Additionally, they were verbally informed about their right to obtain any clinical, psychological and neuroimaging results. Finally, participants were informed about further publications which would result from the project, with the guarantee of anonymization and control of the privacy of their personal information.

### Recruitment and Exclusion Criteria

The recruitment procedure is summarized in table II and subsequently detailed.

**Table II:** General procedure for recruitment.

|  | Activity | Sample size |
| --- | --- | --- |
| Stage 1 | Selection of one municipality which with the closest match in gender, age, ethnic characteristics to of the general Cuban population. This was also a municipality with a large proportion of persons originally from other provinces according the MINSAP and Statistics National Office. La Lisa municipality was selected. A random sample of 30,000 participants from the total population was selected based on the ID card serial numbers. | 30,000 |
| Stage 2 | From the 30,000 participants selected in the previous stage a random subsample of 2019 was selected stratified by age, sex, ethnic origin, educational level, and socio-economic status. | 2019 |
| Stage 3 | The Doctor and nurses of the Family evaluated the willingness to participate and health condition of the candidates from the subsample (n=2019 ). In detail:<br>1) The Family nurses visited each household to provide both printed and verbal information about the project objectives and enquired about preliminary about willingness to participate in the study.<br>2) Those willing to participate went to their local Polyclinic center and for further explanation and signed the Informed Consent.<br>3) The Health Questionnaire was then applied by the nurse in order to carry out a further screening of pathology, cognitive complaints, use of pharmaceutical agents, heavy smoking etc.<br>4) According to the exclusion criteria (see Table III) a further selection of the subsample was carried out for the next stage.<br>Participants with health problems were remained in the overall study, and continue to be followed up to this day with further clinical evaluation and treatment. However, they were excluded from the normative part of the project. | 1439 |
| Stage 4 | At the local Polyclinic center psychiatrist/neurologist applied the:<br>• A standardized and structured psychiatry tool, the MINI International Interview (MINI)<br>• MINIMENTAL State Evaluation (MMSE)<br>• The Edinburg Handedness Questionnaire to classify dexterity<br>A psychometrist applied the: | 530 |
|  | • The Weschler Adult Intelligence Scale (WAIS) III |  |
| Stage 5 | At the local Polyclinic center, blood sample for genetic studies were extracted and additional measures: the blood pressure, body temperature, frequency and heart rate, body weight, height (data not included in this version of the database).<br>Digital EEG recording was the final step at the designated Policlinics | 530 |
| Stage 6 | MRI collection was carried at the CIMEQ Hospital followed by MRI evaluation by a neuroradiologist. | 394 |

#### Stage 1

In agreement with Ministry of Public Health (MINSAP), and due to logistical constraints, a single municipality of the province of Havana City selected for the study, together with the whole structure of the Family Doctor and Polyclinic Centers. Towards this end a study was carried out with a committee from MINSAP and the National Office for Population Studies, to assess the distribution of the following variables of all inhabitants in every municipality in the Province of Habana City: ethnicity, sex, province of origin, and socio-economic status. On the basis of these distributions, the Municipality of La Lisa, was selected for the study since it had the closest match to the general Cuban population. A sample of 30,000 inhabitants in this region was randomly selected from the National Identity Card registry.

#### Stage 2

From the original roster of 30,000, a random subsample of n=2019 was then selected for further processing, being stratified by age, gender, socio-economic status.

Family Doctors (**Stage 3**) then examined the participants records to exclude persons whom they already had ascertained to have health issues. All the remaining participants were visited by the Family nurses who left a printed description of the project and gave a detailed verbal explanation of its aims. As usual for population studies in Cuba, it was explained that there would be no payment for the study, but if a participant needed be absent from his workplace, the local government guaranteed this as a fully paid day. They were also informed about all data acquisition protocols as well as safety measures with a special focus on MRI acquisition and safety. To a great degree, the success of this project was due to the close contact of the Family Doctor and Nurse with the local population, as well as the abundant information provided from the media to the general public about the Cuban Neuroscience Center and its project. This explains a 93% initial willingness to participate in the project. For those participants that gave their written consent, a health questionnaire was applied for further screening and consequently, 580 persons were excluded at this stage from the normative study. In this, as well as in subsequent stages, all participants that did not continue in the normative study followed a separate workflow to ensure specialized diagnostic and intervention by units of the health system, with the same protocol as those continuing in the study. The exclusion criteria used for this stage are listed in Table III. The most prevalent health conditions to exclude participants were diagnosed metabolic syndrome, psychiatric conditions, personal history of severe illnesses, and sensory and motor disabilities.

**Table III:** Exclusion Criteria.

| Criteria | Description |
| --- | --- |
| Medical conditions | Malignant systemic disease requiring chemotherapy, diabetes, thyroid dysfunction (hyper or hypothyroidism), rheumatic disease, muscular dystrophies, liver cirrhosis, sickle cell anemia, Wilson's disease, lupus erythematosus (SLE), malnutrition, cardiovascular diseases, arterial hypertension, infectious diseases, AIDS, respiratory diseases, history of reactions to medications with hypersensitivity type I, current pregnancy or lactation, permanent metal appliances, cardiac pacemakers or other type of metal in any part of the body (fixed prosthesis, fragments of bullet, etc.) |
| Neurological disease | Chronic systemic diseases of the Central Nervous System (CNS), seizures or other attacks, malignant expansive processes of the SNC-radiotherapy, more than one loss of consciousness, cerebrovascular accidents, transient cerebral ischemia, migraine or frequent headaches, severe neuropathies, history of cranial trauma with or without loss of consciousness. |
| Psychiatric Disorder | History of previous psychiatric treatments, tics, stuttering, intellectual disability, suicide attempt, depression, sociopathic behavior, anxiety disorders, panic attacks, schizophrenia, obsessive-compulsive disorder, drug use and abuse, alcoholism, hyperactivity and attention deficit, dementia. |
| Prenatal and perinatal antecedents | Difficult pregnancy as indicated by the obstetrician, preterm delivery, gestational hypertension, diabetes, obesity, early membrane rupture in pregnancy, placenta previa, retroplacental hematoma, other infections. |
| Sleep disorders | Nocturnal terrors, somnambulism, and others. |
| Familiar pathological background | Epilepsy, neurodegenerative diseases, multiple sclerosis, Wilson's disease, schizophrenia, manic depressive disorder. |
| Drug addiction | Alcohol abuse according to DSM-IV definition, non-legal drugs, smoker of more than 12 of cigarettes per day, drugs affecting the CNS, direct and chronic exposure to toxic substances such as pesticides, heavy metals, phosphorous and organic solvents. |
| Neurological physical examination | Any abnormality in the neurological physical examination (hypertonia, hypotonia, asymmetry of reflexes, decrease in visual acuity, nystagmus, etc.). |

In **Stage 4**, participants continuing the study were examined at the polyclinic by specialists in Neurology and Psychiatry, in order to rule out chronic diseases (e.g. addictions, including heavy smoking) or any disorders of the nervous system that would invalidate their participation in the study. Neurological examination was performed following the procedure described in the guidelines published by the U.S. Department of Health and Human Services in 1997 ^15^ and the Mini Mental State Examination^16^ (MMSE) for global cognitive screening. The Mini-International Psychiatric Interview was used for psychiatric evaluation^17^ (Spanish version) and the Weschler Adult Intelligence Scale (WAIS) III for intelligence.

In **Stage 5**, EEG recordings were carried out at the polyclinic. Other measures were obtained the same day such as anthropometric (weight) and blood pressure. Blood samples were extracted, following a join protocol from CNEURO with the National Center for Medical Genetics https://www.ecured.cu/Centro_Nacional_de_Gen%C3%A9tica_M%C3%A9dica for further genetic studies of the Cuban population (to be published separately).

During the stage 4 and 5 specific exclusion criteria (see Table IV).

**Table IV:**
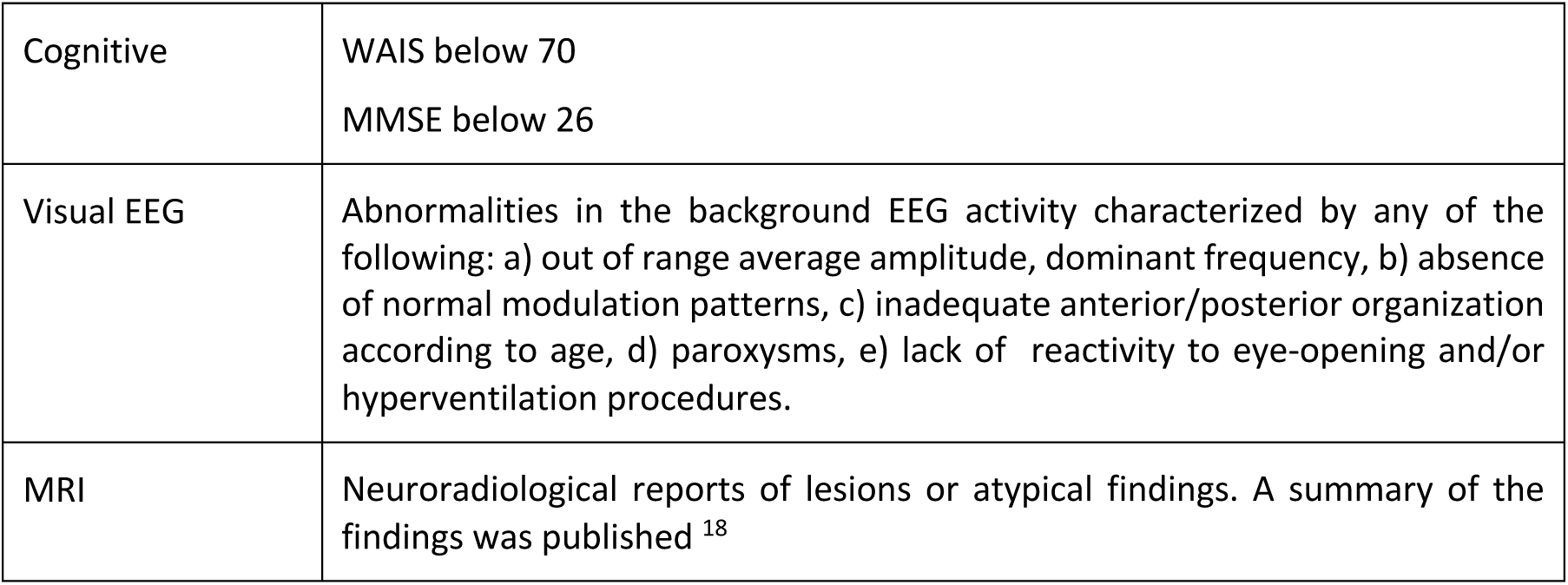
The exclusion criteria according to cognitive, EEG and MRI examination.

|  |  |
| --- | --- |
| Cognitive | WAIS below 70<br>MMSE below 26 |
| Visual EEG | Abnormalities in the background EEG activity characterized by any of the following: a) out of range average amplitude, dominant frequency, b) absence of normal modulation patterns, c) inadequate anterior/posterior organization according to age, d) paroxysms, e) lack of reactivity to eye-opening and/or hyperventilation procedures. |
| MRI | Neuroradiological reports of lesions or atypical findings. A summary of the findings was published <sup>18</sup> |

In **Stage 6**, the MRI studies were carried out at the Center for Medical and Surgical Research la Habana Cuba (in Spanish: Centro Investigaciones Medico-Quirurgicas, CIMEQ https://www.ecured.cu/Centro_de_Investigaciones_M%C3%A9dico_Quir%C3%BArgicas_(CIMEQ)

Additionally, a psychiatrist/psychometrist applied the computerized Reaction Time test at the end of the study for a subsample n=56 of the final sample.

The participants who presented hypertension during this research study were included in a separate study and underwent more specific evaluations such as carotid flow, white matter hyper intensities, eye fund, optic and blood vessel impairments and a set of extra measurements. The analysis of the results of this hypertension study was partially published in ^3^.

## Procedure

### Workflow

Examinations for final participants (Stages 4-6) were carried out in a five-day schedule. You can see the figure 1 below.

**Figure 1.**
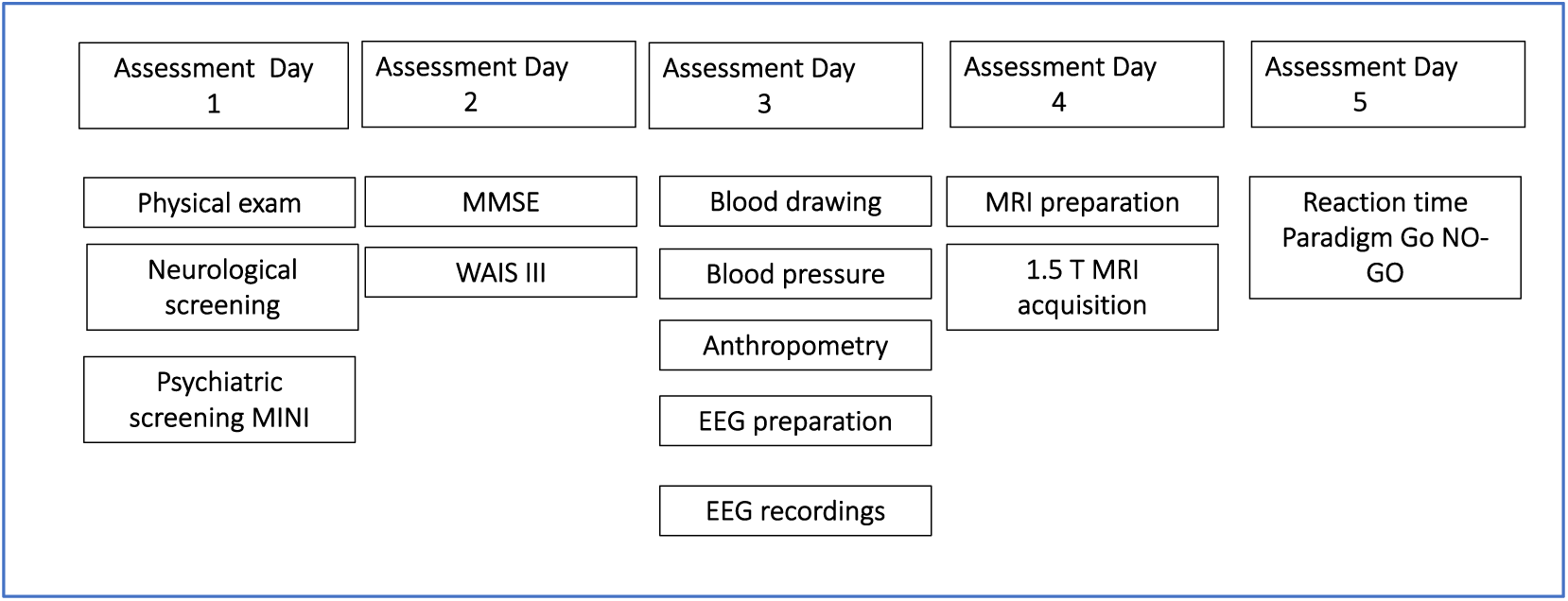
Flow-chart of assessment.

Finally, the sample resulting for this study (N=282) included all the participants who completed all the requisites, after all the steps. The participants included in each measurement and the conjunction between modalities in table V.

**Table V.**

| Measurement | Number |
| --- | --- |
| EEG | 250 |
| MRI | 203 |
| WAIS | 167 |
| MMSE | 156 |
| TR | 56 |
| EEG+MRI | 171 |
| EEG+MRI+WAIS | 142 |
| EEG+MRI+MMSE | 132 |

### Psychological tests

#### MMSE

The Mini Mental State Examination MMSE^16^ is a quick and easy measure of cognitive functioning that has been widely used in clinical evaluation and research involving patients with dementia. In our study was employed as a screening test to exclude participants with cognitive impairment.

#### Intelligence test

Intelligence was assessed using the fully validated and translated to Spanish language version of the David Wechsler Adult Intelligence Scale WAIS-III printed and distributed in Mexico by El Manual Moderno (https://www.worldcat.org/title/wais-iii-escalaweschler-de-inteligencia-para-adultos-iii/oclc/54053545). This scale provided scores for a Full-Scale IQ (FSIQ), Verbal IQ (VIQ), and Performance IQ (PIQ) along with four secondary indices: PO, PS, VC, and WM. The subtests included in each index were as follows: PO: picture completion, block design, matrix reasoning; PS: digit-symbol coding and symbol search; VC: vocabulary, similarities, information, comprehension; and WM: arithmetic, digit span, letter-number sequencing.

The intelligence raw measures were scored according to the official normative data included in the printed version of WAIS-III. However, to avoid cultural bias, they were subsequently standardized with information from the Cuban sample to produce scores of the specific performance, adjusted for age for our population. The results about how the white matter (FA-tracts based) predicts fluid and crystalized intelligence has been published using this dataset^19^.

#### Go No Go test

For a sub-set of 56 participants, reaction times were recorded using a go-no go paradigm which consisted in a visual attention task, implemented using the psychophysiology software for cognitive stimulation Mindtracer^20^ (N_P-SW 1.3 v.2.1.0.0 Neuronic S.A.)

The task consisted of 500 trials, 25% in GO condition and 75% in NO-GO condition, where the stimuli was a set of letters: P B X E A S. The instructions for the participants was: Simple Reaction Time (RT1) consign: “Press the space bar when “S” appears at the screen”. Complex Reaction time (RT2) consign: “Press the space bar only when the letter “S” appears preceded by a letter “A”. The two tasks were presented to all the participants in the same consecutive order. For a description of the facilities of the software see figure 2.

**Figure 2.**
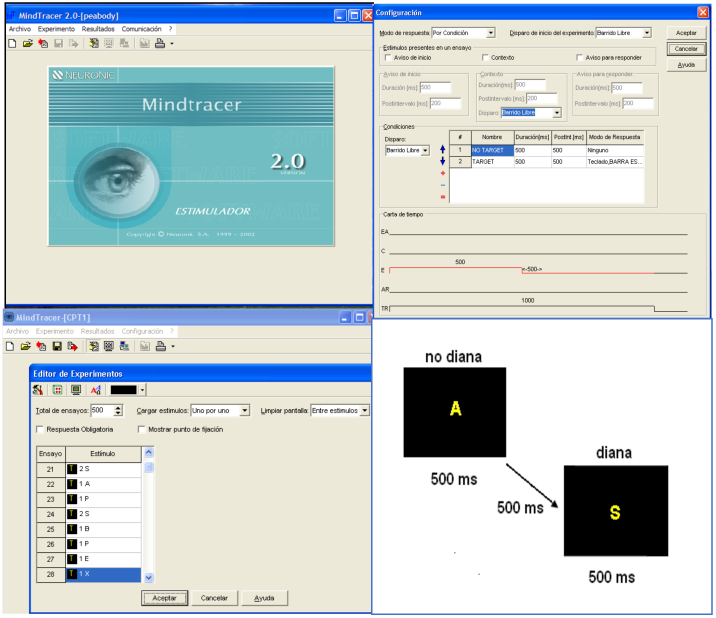
Top left.- Main display of the software MINDTRACER for the preparation and presentation of the stimuli of the reaction time task. Top right.- Software options to design the experimental task. Bottom left.- Description of the different trials. Bottom right.- Design of the presentation of the complex stimuli.

### EEG recordings

Resting state EEG was recorded using the digital electroencephalograph system (MEDICID 5-64 and MEDICID 5-128) (www.neuronic.cu) with differential amplifiers and gain of 10,000. Electrodes were placed according to the 10–10 International System with a customized electrode cap. Linked earlobes were used as the EEG reference. Electrode impedances were considered acceptable if less than 5 KΩ. The band pass filters parameters was 0.5–50 Hz and 60 Hz notch, and sampling period of 200 Hz. The EEG was recorded in a temperature and noise-controlled room while the participant was sitting in a reclined chair. All individuals were asked to relax and remain at rest during the test to minimize artifacts produced by movements, and to avoid excessive blinking. The participants received instructions to have enough sleep the previous night, take breakfast and wash the hair before attending this appointment. See table VI for a summary of the technical parameters of the EEG. The structure of raw EEG recording was generated in the default format of the MEDICID neurometrics system (*.plg extension), which later are converted to standard BIDS format. See data records section.

**Table VI:**

|  |  |
| --- | --- |
| Equipment | MEDICID 5 |
| Number of channels | 64/128 |
| Gain | 20,000/1000 |
| Impedances | <10 KΩ. |
| Line Frequency | 60 Hz |
| Amplifiers bandwidth | 0.5 to 50 Hz |
| Sampling frequency | 5msec./<br>200Hz. |
| Room Temperature | 20 to 22<sup>0</sup> C. |

### Electrode placement

Two different montages were employed, one with 64 channels and other with 120 channels as illustrated in Figure 3 with different colors black (64) and white (120) to identify each montage. The nomenclature of the electrodes employed in the MEDICID system and their standardization is included in supplementary material 1 at the end of this document.

**Figure 3.**
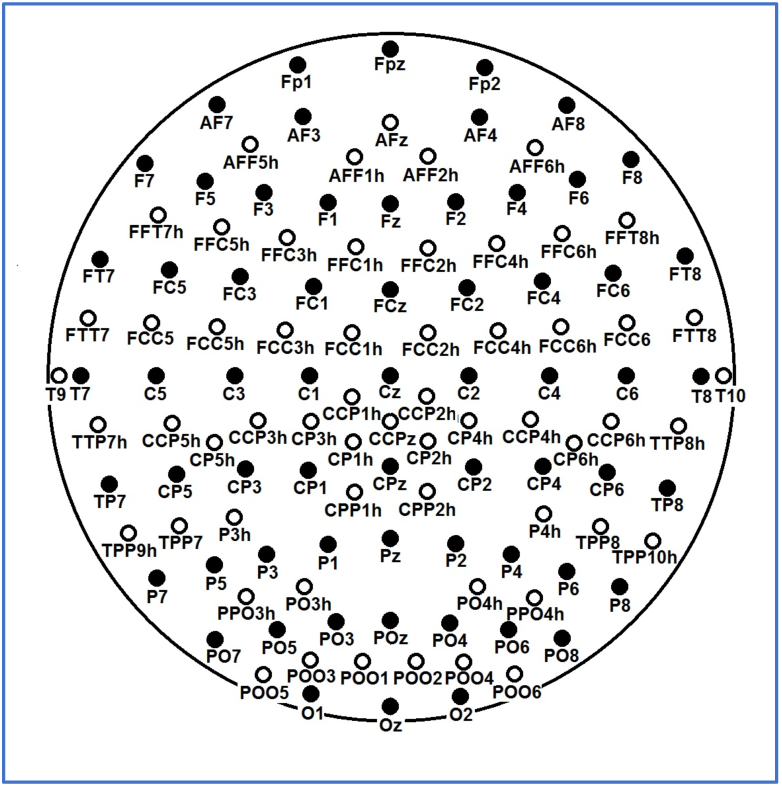
Schematic representation of the electrodes position. In black the sub-set of electrodes employed in the 64 channels (maximum 62 cephalic electrodes). In white the configuration for the 120 channels (maximum 120 cephalic electrodes). In both configuration, three channels are employed to record the Electro-oculogram and Electrocardiogram.

#### Description of the EEG protocol comprising the following participant condition

1. Baseline: resting state EEG with closed Eyes (state A), 10 minutes
2. Reactivity: this test consisted in the consecutive opening and closing eyes with an interval of 12 seconds. Open eyes (state B), 5 minutes, where the participant was instructed to look at a point, keeping the pupils fixed.
3. Hyperventilation (HPV): Dividing it in the first minute HPV1 (state C), the subject was instructed to start taking air through the nose and to breathe deeply. The second minute HPV2 (state D), and HPV3 (state E), this last minute less deep and more frequent. Total 3 minutes.
4. Recovery (state F): The last step is the recovery of the patient after the HPV, which lasted around one and half minute, but were recorded for 2 full minutes.

Note that the subject’s recordings were monitored continuously by the technician, in order to avoid contamination of the EEG with the electromyogram interference, other changes in the direct current level due to sweating, and also to prevent drowsiness. Any of this contamination were annotated online by the technician.

Therefore, recordings of at least half an hour were ensured. A design requirement was to have enough valid EEG to carry subsequent frequency domain analysis. For this EEG epochs for are necessary, each consisting of 256 time samples, or 2.56 seconds being marked on line continuous EEG recordings. Due to the high density of electrodes, the number of epochs for further analysis was guaranteed to be at least 50 windows for 64 channels and 80 windows for 120 channels (For details on analysis see ^10,12^.

### MRI procedure

MRI: Magnetic resonance imaging (MRI) was performed on a 1.5 Tesla scanner (MAGNETOM Symphony Siemens Erlangen Germany). Over the course of MRI data acquisition, the scanner remained stable and did not undergo any major maintenance or updates which would systematically affect the quality of data provided here. The total measurement time was 45 minutes. See the MRI protocol used in Table VII.

**Table VI:** MRI Protocol

| Protocol | Description |
| --- | --- |
| T1w | • Patient position: First Supine Head<br>• Sequence: 3D MPRAGE<br>• FOV: 256 x 256 x 160 mm<br>• Voxel dimensions: 1 x 1 x 1 mm<br>• Inversion Time (TI): 1100 ms<br>• Repetition Time (TR): 3000 ms<br>• Echo Time (TE): 3.93 ms<br>• Flip Angle: 15 degrees<br>• Series Name: t13d_anatVOL<br>• Name of the sequence: tfl3d1_ns brusque and remarkable. |
| Field Maps | • Sequence: 2D Double-echo gradient echo field map<br>• FOV: 224 x 224 mm<br>• Imaging matrix: 64 x 64<br>• In-plane resolution: 3.5 x 3.5 mm<br>• Number of slices: 40<br>• Slice orientation: Axial<br>• Slice thickness: 3.5mm<br>• Slice gap: 0 mm<br>• Repetition Time (TR): 672ms<br>• First Echo Time (TE1): 7.71 ms<br>• Second Echo Time (TE2): 12.47 ms<br>• Acquisitions: 1<br>• Flip Angle: 60 degrees<br>• Image type: Magnitude and Phase<br>• Name of the Series: fieldmap pha or fieldmap_mag depending on the type<br>• Name of the Sequence: fm2d2r. |
| Diffusion weighted images (DWI) Variant 1 | • Sequence: 2D Single-Shot Gradient Echo Echo-Planar-Imaging (GE-EPI)<br>• Parallel acquisition: none<br>• Multiband: none<br>• FOV: 256x256 mm<br>• Imaging matrix: 128 x 128<br>• Phase-Encoding direction: AP<br>• In-plane resolution: 2 x 2 mm<br>• Number of slices: 25<br>• Slice orientation: Axial<br>• Slice thickness: 3 mm<br>• Slice gap: 3 mm<br>• Repetition Time (TR): 7000 ms<br>• Echo Time (TE): 160 ms<br>• Flip Angle: 90 degrees<br>• Acquisitions: 6<br>• Diffusion gradient scheme: Siemens MDDW<br>• Number of diffusion gradient directions: 12<br>• Echo Spacing: 1.149 ms<br>• Readout Time: 145.92 ms<br>• Image type: Magnitude<br>• Series Name: ep2d_diff12dir<br>• Name of the Sequences: ep_b0, ep_b1200 # 0 ... 11<br>This DWI sequence was repeated in a second run with the same parameters. The only difference was the position of the slices. They were translated parallel to the axis normal to the slice (axial plane) so that the single from the gaps of the first run were acquired in the second run. Consequently, the gaps of the second run occupied the regions from which the signal in the first run was acquired. In this way the entire brain was covered with a total of 50 slices. |
| Diffusion weighted images (DWI) Variant 2 | • Sequence: 2D Single-Shot Gradient Echo Echo-Planar-Imaging (GE-EPI)<br>• Parallel acquisition: none<br>• Multiband: none<br>• FOV: 256x256 mm<br>• Imaging matrix: 128 x 128<br>• Phase-Encoding direction: AP<br>• In-plane resolution: 2 x 2 mm<br>• Number of slices: 25<br>• Slice orientation: Axial<br>• Slice thickness: 3 mm |
|  | • Slice gap: 0 mm<br>• Repetition Time (TR): 7000 ms<br>• Echo Time (TE): 160 ms<br>• Flip Angle: 90 degrees<br>• Acquisitions: 6<br>• Diffusion gradient scheme: Siemens MDDW<br>• Number of diffusion gradient directions: 12<br>• Echo Spacing: 1.41 ms<br>• Image type: Magnitude<br>This sequence was not repeated. |

Table VII summarizes the technical specifications for the acquisition of the MRI. Note that only 203 participants have useful MRI images. With respect to diffusion images, we collected 201 participants with two variants: Variant 1 was performed on 148 subjects and Variant 2 on 53 subjects. The DWI specific parameters can be found inside the BIDS format of each participant. All DWIs were visually inspected and those which presented either technical or pathological defects were discarded.

Different analysis and type of processing using this MRI dataset already has been published. One study demonstrated how the surface area could explain the morphological connectivity of brain networks ^21^ (Sanabria Neuroimage 2010). Other study explained the substantial inter-individual variability on the neuroanatomical determinants of EEG spectral properties using the DWI-fractional anisotropy ^22^ (Valdes-Hernandez 2010 Neuroimage). Two papers studied the human brain anatomical network via diffusion-weighted MRI and Graph Theory characterizing brain anatomical connections ^23,24^ (Iturria 2007,2008 Neuroimage). Another paper presented a general framework for tensor analysis of single-modality model inversion and multimodal data fusion using our neuroimaging data as an example^25^ (Karahan Tensor IEEE 2015).

## Code Availability

The following in-house Windows software developed by EAV will be soon available at GITHUB.

1. EEG-Anonymizer: to erase all the personal information stored in the EEG recordings, which could facilitate the identification of the participants.
2. PLG2BIDS: to read the original EEG recordings in NEURONIC format and convert them to BIDS structure.
3. Combine EEG-MRI-BIDS: to combine BIDS-EEG and MRI-BIDS into only one structure.
4. Unwrap: As part of the MRI quality control process, several MRI T1images studies were fixed when a wrap-around artifact (without overlapping on head) was detected.
5. Quality Control MRI: Automatic inspection was performed to check the protocol parameters of the DWI images and generate a file with the value of the parameters.

## Data Record

BIDS (Brain Imaging Data Structure) is the new standard for the organization and description of the datasets containing neuroimaging (MRI, MEG, EEG, iEEG, NIRS, PET) and behavioral information ^26^.

Based in this BIDS structure, we developed a methodology with the following steps:

1. Anonymization of EEG recordings and MRI scans.
2. Defacing of the MRI scans
3. Conversion of EEG recordings and MRI to BIDS-EEG
4. Validation of the BIDS structure

### Anonymization

We developed an application (EEG-Anonymizer) to erase all the personal information stored in the EEG recordings, which could facilitate the identification of the participants. This application generated a security copy of the personal information before its elimination.

The anonymization of the MRI neuroimages was performed using the script Dicat.py V 1.2 (https://github.com/aces/DICAT), https://github.com/aces/DICAT developed by MCIN (McGill Center for Integrative Neuroscience) Montreal, Canada. All the data was anonymized with DICAT in 4 folders (ID, Patient_Name, Sex, Birth_Date).

### Defacing

The defacing process consisted in the elimination of the section with the face of the subject inside the anatomical MRI. This prevent the identification of the subject if a posterior 3D rendering is employed with the MRI scan. The software employed was the Mri_deface V 1.2, del FreeSurfer https://surfer.nmr.mgh.harvard.edu/fswiki/mri_deface

### Conversion to BIDS

#### EEG

We developed an ad-hoc application (PLG2BIDS:) to read the original EEG recordings in NEURONIC format and convert them in BIDS structure. This application is designed to read either individual EEG recordings or folders with multiple recordings and is able to update a current BIDS structure with new recordings.

#### MRI

The conversion of MRI neuroimages to BIDS structure was using the Dcm2Bids https://github.com/cbedetti/Dcm2Bids which generate the MRI BIDS structure with the original data in format DICOM.

The BIDS-EEG and MRI-BIDS structures were combined in only one structure using a software Combine EEG-MRI-BIDS.

### Validation

The final step was the validation of the BIDS structure using the web bids-validator. https://bids-standard.github.io/bids-validator/

All the dataset was also imported in Longitudinal Online Research and Imaging System LORIS)v20.2. https://mcin.ca/technology/loris/

## Technical Validation

For the quality control of the EEG, MRI and psychological tests data of this study was implement in a workflow:

### EEG

1. The ocular movements and other incidences were annotated by the technician during the recordings.
2. A Board Certified clinical neurophysiologist reviewed the recorded raw EEG by visual inspection to provide:
  a. Overall assessment of the EEG recording which if of insufficient quality could lead to a repetition of the EEG recording.
  b. Scoring of semi-quantitative scales for abnormalities which could motivated the exclusion of the participant from the normative sample.
  c. Selection of artifact free EEG segments useful for further analysis in the continuous recordings.

### MRI

1. Automatic inspection was performed to check the protocol parameters of the MRI images using in-house software which generate a file with the value of the parameters.
2. Visual inspection by several clinical radiologists to detect abnormalities to decide if the participant should be excluded.
3. As part of the MRI quality control process, several MRI T1 images studies were fixed when a wrap-around artifact (without overlapping with the head) was detected by using the in-house Quality Control MRI app.

### Psychological tests and behavioral information

1. Supervision of the assessment sessions by one board certified Clinical Neuropsychologist.
2. After input to the system, curation of the clinical, psychological and demographical data was carried by a CNEURO team, assisted by statistical summary tools, to ensure quality control.

## Usage Notes

Each participant was assigned one ID with the structure **CBMxxxx** 3 characters identifying “Cuban Brain Mapping” and four digits indicating the number or order of the participant in the dataset. Note that code is the same to the different data modalities of the subject.

The EEG-BIDS, the BIDS-MRI and psychological, demographic and clinical data in a standard ASCII (∗.csv, ∗.txt) will be available from https://cloud2.cneuro.cu.

All the datasets have been also stored in the McGill Centre for Integrative Neuroscience (MCIN) network. The dataset will be available by request at https://cuba-open-dev.loris.ca/

More details about the access to the dataset would be published in the final version of this paper.

## Acknowledgements

The second wave of the CHBMP was carried out during the National Program of Disabilities initiated by the Ministry of Public Health in 2004 and directed by Vice-Minister Marcia Cobas Ruiz. Our gratitude to all the Cuban participants who volunteered to participate. Very special thanks to the nurses and the staff of the Policlinic “Elpidio Berovides” and “Aleida Fernandez” of the municipality of La Lisa, La Habana Cuba who were indispensable to carry out this project.

We would like to thank the contributions of principal investigators, researchers, and staff personnel of the Cuban Neuroscience Center and former members of the CHBM project who are not authoring this paper, but were founders: Pedro A. Valdes Hernandez, Yasser Iturria, Lester Melie-Garcia and Gertrudis Hernandez Gonzalez. This project was carried out in collaboration with the National Center for Clinic Genetic directed by Beatriz Marcheco.

The authors would like to thank the support from the National Natural Science Foundation of China (NSFC) projects (81861128001, 61871105, 61673090, and 81330032) and the CNS Program of UESTC (No. Y0301902610100201). This work is part of the Cuba China Canada (CCC) axis platform of neuroinformatics.

## Author Contributions

PVS was the principal investigator of this project. PVS and MVS with JBB and LGG conceived, this project, directed the data collection and wrote the first version of this article. MLB designed and directed the neuropsychological assessment and the analysis of cognitive data. IRG and EAV were in charge of the cooperation to upload the dataset in the MCIN inside LORIS. EAV did the processing and visualization of neuroimaging data; TVA was in charge of the protocols, recordings and visual inspection of the EEG; LVU was in charge of logistic and organization of the study. ACE gave the strategic and institutional support of MCIN, and SD, CM, ZM, LCM, CR and SB were the Canadian team in charge of the CBRAIN platform and LORIS software integration of the CHBMP data. PVS, LGG, JBB and MLB wrote the final version of the paper.

## Supplementary material 1

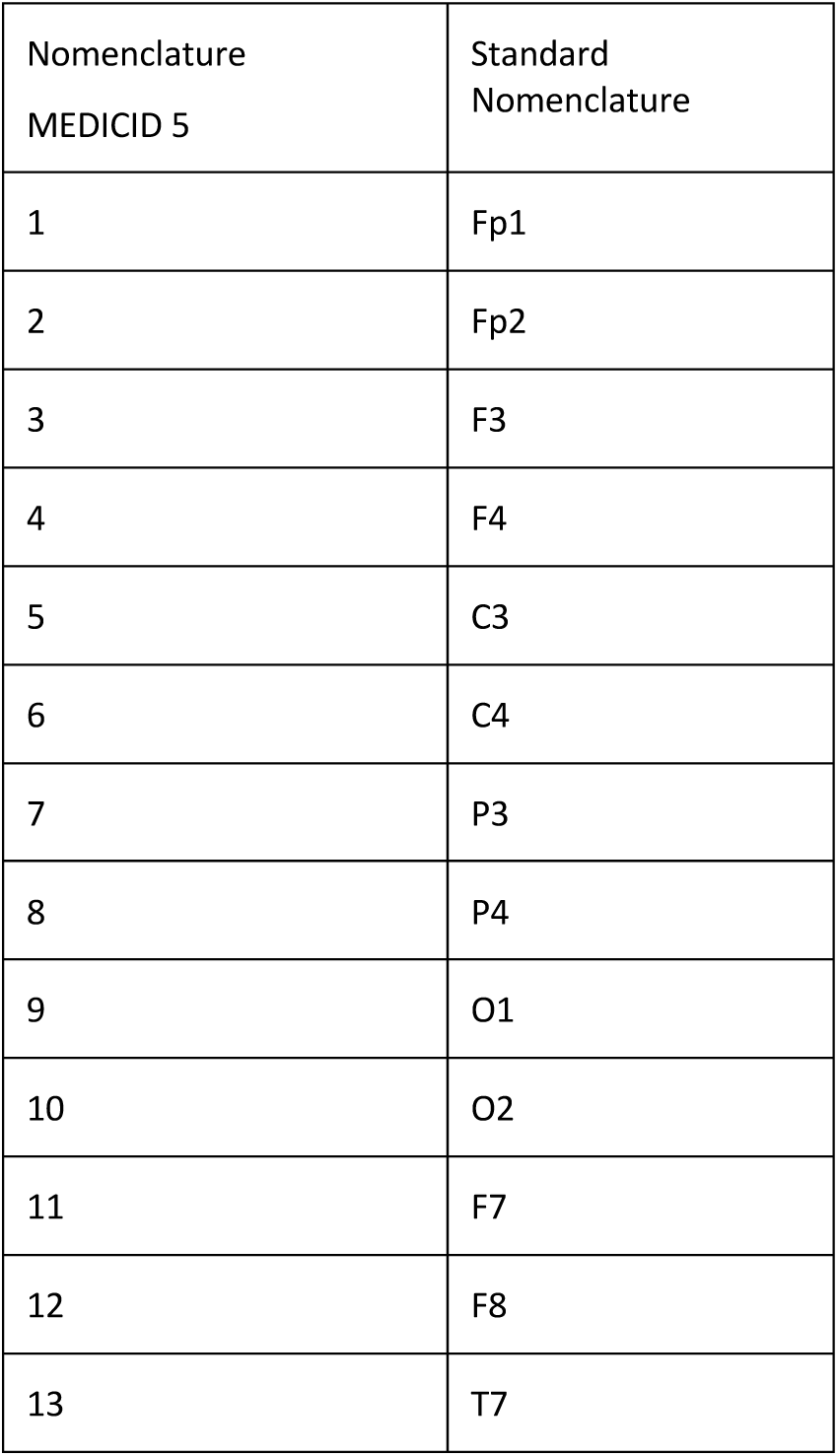

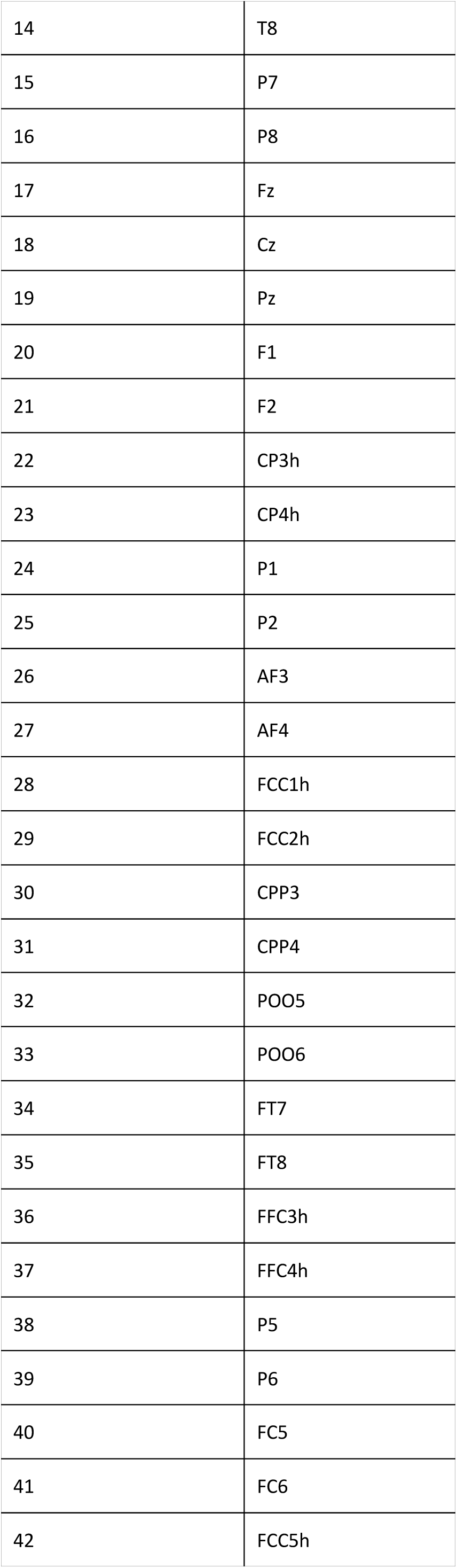

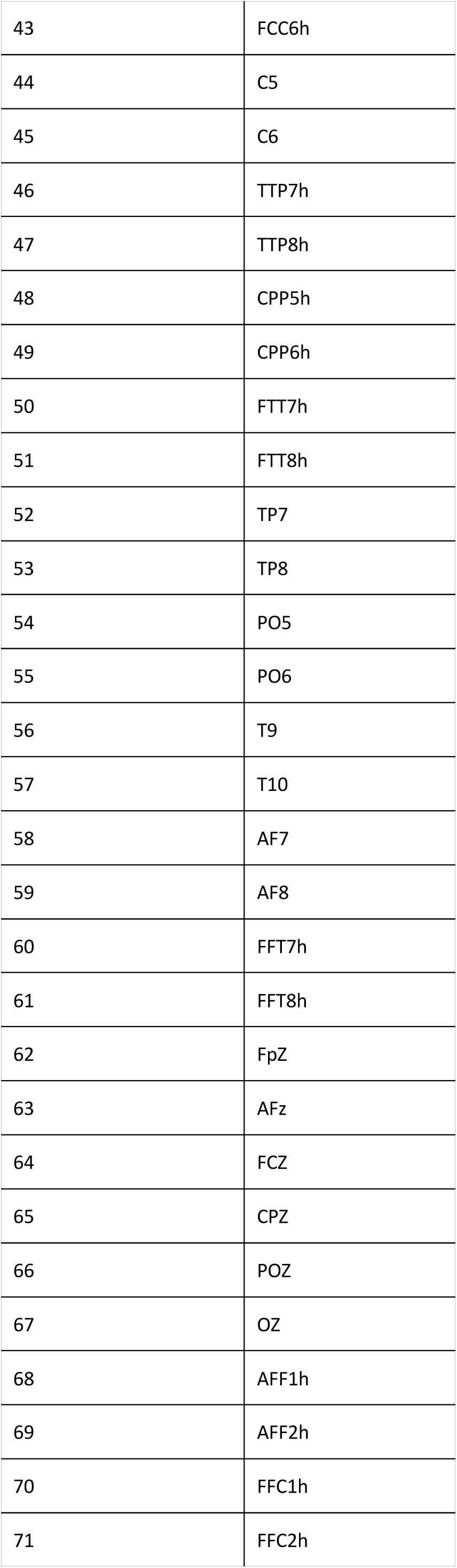

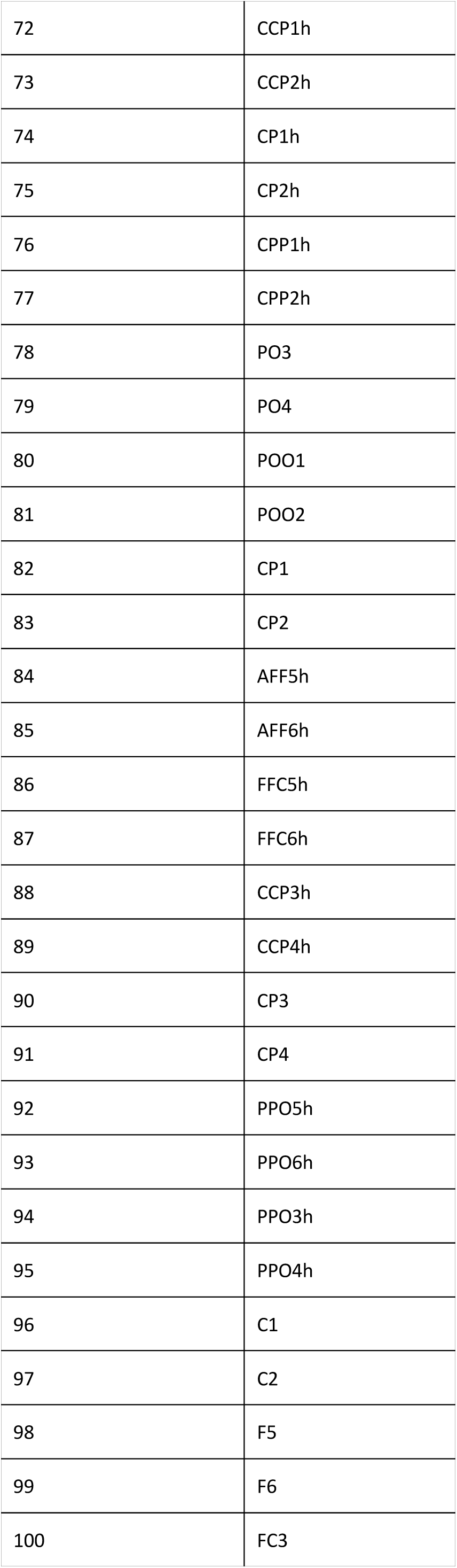

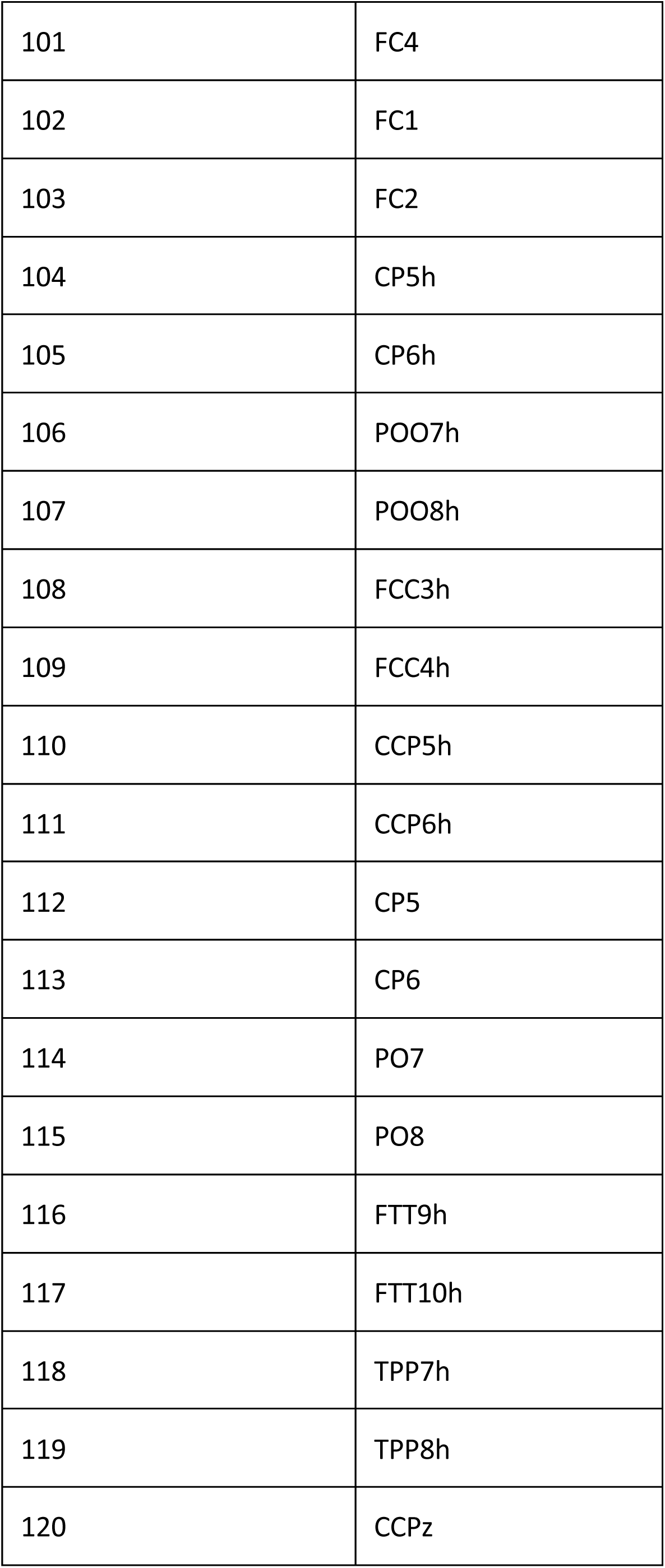

## References

1. Babayan, A. et al. Data descriptor: A mind-brain-body dataset of MRI, EEG, cognition, emotion, and peripheral physiology in young and old adults. Sci. Data 6, 1–21 (2019).

2. Alexander, L. M. et al. Data Descriptor: An open resource for transdiagnostic research in pediatric mental health and learning disorders. Sci. Data 4, 1–26 (2017).

3. Hernandez-Gonzalez, G. et al. Multimodal quantitative neuroimaging databases and methods: the Cuban Human Brain Mapping Project. Clin. EEG Neurosci. 42, 149–159 (2011).

4. Das, S., Zijdenbos, A. P., Harlap, J., Vins, D. & Evans, A. C. LORIS: a web-based data management system for multi-center studies. Front. Neuroinform. 5, 1–11 (2012).

5. John, E. R. et al. Neurometrics. Science (80-.). 196, 1393–1410 (1977).

6. Valdés, P. et al. QEEG in a Public Health system. Brain Topogr. 4, 259–266 (1992).

7. Szava, S. et al. High resolution quantitative EEG analysis. Brain Topogr. 6, 211–219 (1994).

8. Valdés, P. et al. Frequency domain models of the EEG. Brain Topogr. 4, 309–19 (1992).

9. Evans, A. et al. 3D statistical neuroanatomical models from 305 MRI volumes. in Proceedings of IEEE-Nuclear Science Symposium and Medical Imaging Conference 1813–1817 (1993).

10. Bosch-Bayard, J. et al. 3D statistical parametric mapping of EEG source spectra by means of variable resolution electromagnetic tomography (VARETA). Clin. Electroencephalogr. 32, 47–61 (2001).

11. Bosch-Bayard, J., Galan-Garcia, L., Aubert-Vázquez, E., Virues-Alba, T. & Valdes-Sosa, P. A. Resting state healthy EEG: the first wave of the Cuban normative database. Front. Neurosci. submitted, (2020).

12. Bosch-Bayard, J. et al. A quantitative EEG toolbox for the MNI Neuroinformatics ecosystem: normative SPM of EEG source spectra. Front. Neuroinform. (2020).

13. Park, H. J. et al. Statistical parametric mapping of LORETA using high density EEG and individual MRI: Application to mismatch negativities in schizophrenia. Hum. Brain Mapp. 17, 168–178 (2002).

14. World Medical Association. World Medical Association Declaration of Helsinki. Ethical Principles for Medical Research Involving Human Subjects. Bull. World Health Organ. 79, 373–374 (2001).

15. Services, U. S. D. of H. and H. Neurological Single Examination. 1 (1997).

16. Folstein, M., Robins, N. & Helzer, J. The mini-mental state examination. Arch. Gen. Psychiatry 40, 812 (1983).

17. Lecrubier, Y. et al. La Entrevista Neuropsiquiátrica Internacional Reducida (MINI). Una entrevista diagnóstica estructurada breve: fiabilidad y validez según la CIDI. Eur. Psychiatry Spanish Ed. 5, 13–21 (1998).

18. Hernández-gonzález, G. D. L. Á., Álvarez-sánchez, M. & Jordán-gonzález, J. Prevalencia de hallazgos incidentales en las imágenes de Resonancia Magnética: Proyecto Cubano de Mapeo Cerebral Humano. Rev. CENIC Ciencias Biológicas 43, 1–6 (2012).

19. Góngora, D., Jahanshahi, M., Vega-hernández, M., Valdés-sosa, P. A. & Bringas-vega, M. L. Crystallized and fluid intelligence are predicted by microstructure of specific white-matter tracts. Hum. Brain Mapp. 1–11 (2019). doi:10.1002/hbm.24848

20. Borrego Hernández, M., Díaz-Comas Martínez, L. & Bobes León, M. A. MINDTRACER 2.0, SISTEMA DE ESTIMULACIÓN PARA ESTUDIOS COGNITIVOS. in BioInformatica2007 6. (CD)BIO030).ISBN: 978-959-286-002-5. 1–10 (2007).

21. Sanabria-Diaz, G. et al. Surface area and cortical thickness descriptors reveal different attributes of the structural human brain networks. Neuroimage 50, 1497–510 (2010).

22. Valdés-Hernández, P. a et al. White matter architecture rather than cortical surface area correlates with the EEG alpha rhythm. Neuroimage 49, 2328–39 (2010).

23. Iturria-Medina, Y. et al. Characterizing brain anatomical connections using diffusion weighted MRI and graph theory. Neuroimage 36, 645–660 (2007).

24. Iturria-Medina, Y., Sotero, R. C., Canales-Rodríguez, E. J., Alemán-Gómez, Y. & Melie-García, L. Studying the human brain anatomical network via diffusion-weighted MRI and Graph Theory. Neuroimage 40, 1064–1076 (2008).

25. Karahan, E., Rojas, P. A., Bringas-vega, M. L., Valdes-Hernandez, P. A. & Valdes-Sosa, P. A. Tensor Analysis and Fusion of Multimodal Brain Images. Proceeding IEEE 103, (2015).

26. Pernet, C. R. et al. EEG-BIDS, an extension to the brain imaging data structure for electroencephalography. Sci. data 6, 103 (2019).

